# Ultrafast and interpretable single-cell 3D genome analysis with Fast-Higashi

**DOI:** 10.1101/2022.04.18.488683

**Authors:** Ruochi Zhang, Tianming Zhou, Jian Ma

**Affiliations:** Computational Biology Department, School of Computer Science, Carnegie Mellon University, Pittsburgh, PA 15213, USA

## Abstract

Single-cell Hi-C (scHi-C) technologies can probe three-dimensional (3D) genome structures in single cells and their cell-to-cell variability. However, existing scHi-C analysis methods are hindered by the data quality and the complex 3D genome patterns. The lack of computational scalability and interpretability poses further challenges for large-scale scHi-C analysis. Here, we introduce Fast-Higashi, an ultrafast and interpretable method based on tensor decomposition that can jointly identify cell identities and chromatin meta-interactions. Fast-Higashi is able to simultaneously model multiple tensors with unmatched features of different sizes. A new partial random walk with restart (Partial RWR) algorithm in Fast-Higashi efficiently mitigates data sparseness. Extensive evaluations on real scHi-C datasets demonstrate the advantage of Fast-Higashi over existing methods for embedding, leading to improved delineation of rare cell types and better reconstruction of developmental trajectories. Fast-Higashi can directly infer chromatin meta-interactions, identify 3D genome features that define distinct cell types, and help elucidate cell type-specific connections between genome structure and function. Moreover, Fast-Higashi can be generalized to incorporate other single-cell omics data. Fast-Higashi provides a highly efficient and interpretable scHi-C analysis solution that is applicable to a broad range of biological contexts.

## Introduction

The advent of high-throughput whole-genome mapping methods for the three-dimensional (3D) genome organization such as Hi-C [1] has revealed distinct features of chromatin folding in various scales within the cell nucleus, including A/B compartments [1], subcompartments [2, 3], topologically associating domains (TADs) [4, 5], and chromatin loops [2]. These multiscale 3D genome features collectively contribute to vital genome functions such as transcription [6, 7]. However, the variation of 3D genome features and their functional significance in single cells remain poorly understood [8, 9]. The recent advances of single-cell Hi-C (scHi-C) technologies have provided us with unprecedented opportunities to probe chromatin interactions at single-cell resolution, from a few cells of given cell types [10–13] to thousands of cells from complex tissues [14–16]. These new technologies and datasets have the promise to unveil the connections between genome structure and function in single cells for a wide range of biological contexts in health and disease [9].

However, the complexity of scHi-C data has created enormous analysis challenges. Computational methods HiCRep/MDS [17], scHiCluster [18], LDA [12], and more recent deep learning based methods 3DVI [19] and Higashi [20] have been developed for the embedding and imputation of the sparse scHi-C data. These existing methods, however, cannot (i) effectively infer informative embeddings for the delineation of rare cell types in complex tissues, (ii) directly identify critical chromatin organizations related to cell type-specific genome functions, and (iii) efficiently operate on large-scale datasets with limited memory resources. It remains an open question on how to develop effective computational methods that can identify rare cell types in complex tissues in an interpretable manner with high scalability, key to understanding the interplay among chromatin organization, genome functions, and cellular phenotypes.

The recent scHi-C embedding method scHiCluster [18] uses linear convolution and random walk with restart to impute the sparse contact maps and applies principal component analysis (PCA) on the imputed maps. This requires the storage of all imputed dense maps in the memory, drastically limiting its application to datasets with a large number of cells at high resolution. More recently, deep learning based scHi-C analysis methods have been proposed, including 3DVI [19] based on a deep generative model and our recent work Higashi [20] that uses a new hypergraph neural network architecture [21]. Both methods suggest better embedding results with Higashi being the first scHi-C embedding approach to demonstrate that the complex neuron subtypes in human prefrontal cortex can be revealed by chromatin conformation only. However, due to the computation-intensive nature of neural networks, the scalability of both methods has much room for improvement for large-scale datasets. For 3DVI, individual variational autoencoders are trained for each genomic distance and each chromosome, leading to thousands of deep neural network models to be trained. For Higashi, since the model treats each contact of scHi-C data as individual samples, it takes a long time to fully iterate over the dataset or to train the model till convergence. Crucially, methods for improving the interpretability of the embeddings for scHi-C data are significantly lacking, limiting our understanding of 3D genome structure-function connections for a diverse set of cellular phenotypes.

Here, we develop Fast-Higashi, a new interpretable and scalable framework for embedding and integrative analysis of scHi-C data. We propose a novel concept for single-cell 3D genome analysis, called “meta-interactions” (analogous to the definition of metagenes in scRNA-seq analysis [22]), to improve the model interpretability. Our proposed Fast-Higashi algorithm jointly produces embeddings and meta-interactions for a given scHi-C dataset. Applications to various scHi-C datasets of complex tissues demonstrate that Fast-Higashi has overall comparable or even better embeddings than existing methods but is significantly faster than neural-network based methods (*>*40x faster than 3DVI and *>*9x faster than Higashi), enabling ultrafast delineation of cell subtypes or rare cell types in different biological contexts. Moreover, Fast-Higashi is able to infer critical chromatin meta-interactions that define cell types with strong connections to cell type-specific gene transcription. Fast-Higashi is the fastest and most scalable method for large-scale scHi-C data analysis to date.

## Results

### Overview of Fast-Higashi

**Fig**. 1a illustrates the overall architecture of Fast-Higashi, which is an interpretable model for scHi-C analysis. In Fast-Higashi, scHi-C contact maps from different chromosomes are represented as multiple three-way tensors. Then a tensor decomposition model is utilized and generalized to simultaneously model these 3-way tensors that share only a single dimension (single cells). The tensor decomposition model takes the tensor representation of scHi-C data as input and decomposes the tensors into multiple factor matrices (**Fig**. 1a) to jointly infer cell embeddings as well as meta-interactions. These meta-interactions manifest the aggregated patterns of chromatin interactions, which are analogous to the concept of metagenes in scRNA-seq analysis. Each meta-interaction corresponds directly to a specific dimensions of the cell embeddings, providing a direct solution to interpret the association between embedding results and 3D genome features. We derived the mini-batch optimization procedure for the tensor decomposition model such that it can efficiently model tensors with drastically different sizes and effectively scale to scHi-C datasets with a large number of cells or at high resolutions. To mitigate the sparseness of the scHi-C contact maps while keeping the advantages of mini-batch training, we proposed a novel partial random walk with restart algorithm (Partial RWR, **Fig**. 1b) that efficiently imputes the sparse scHi-C contact maps before passing them to the tensor decomposition model. The detailed description of the tensor decomposition model and the Partial RWR module are described in **Methods**. Additional descriptions of the optimization procedure can be found in **Supplementary Methods**.

**Figure 1:**
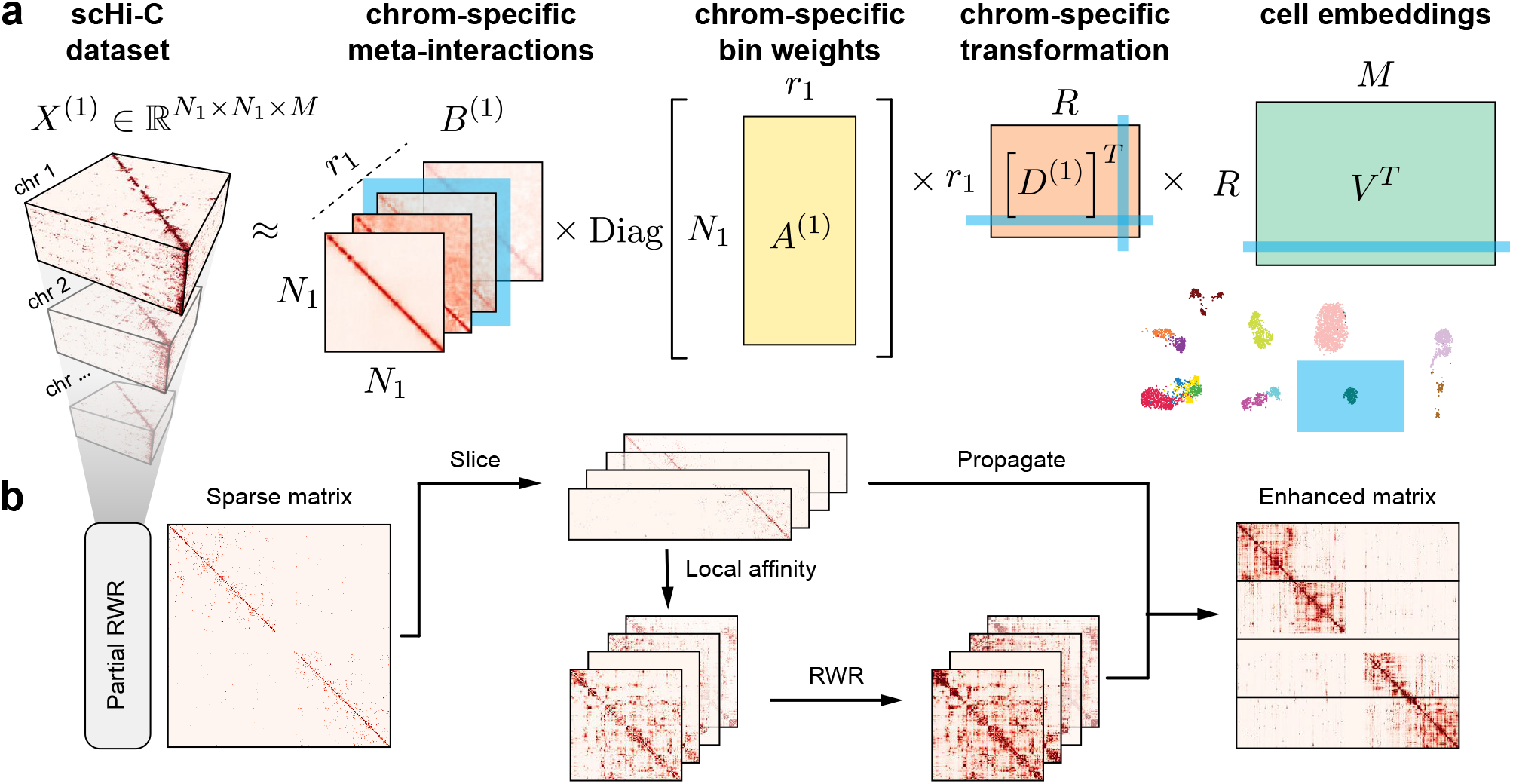
Overview of Fast-Higashi. **a**. Workflow of the Fast-Higashi algorithm. Given an input scHi-C dataset of *k* chromosomes, Fast-Higashi models it as *k* 3-way tensors. The tensor of chromosome *c* is denoted by X^(c)^, where the first two dimensions correspond to genomic bins and the last dimension corresponds to the single cells. Fast-Higashi then decomposes the tensors *X*^(c)^ into four factors: a set of meta-interactions(B^(c)^), a genomic bin weights indicating importance for each bin (A^(c)^), a cell embedding matrix *V* that is shared across all chromosomes, and a chromosome-specific transformation matrix D^(c)^that transforms the shared cell embeddings into chromosome-specific ones. **b**. Workflow of the partial random walk with restart (Partial RWR) algorithm. The Partial RWR is integrated into the Fast-Higashi framework. When calculating the decomposed factors for frontal slices of the tensor *X*^(c)^, the corresponding slices would be imputed through Partial RWR first. The imputation process includes the calculation of local affinity, standard RWR algorithm, and information propagation using both sliced tensor and the RWR imputed affinity matrix.

### Fast-Higashi achieves accurate and fast embedding of scHi-C data

We systematically evaluated the performance of Fast-Higashi for generating embedding vectors for various scHi-C datasets. To demonstrate the effectiveness of Fast-Higashi for delineating subtle cell-to-cell variability of 3D genome features, we applied it to three recent scHi-C datasets of complex tissues at 500Kb resolution. These datasets include the Tan et al. dataset [14], the Lee et al. dataset [15], and the Liu et al. dataset [16] (see **Supplementary Methods** A.1 for data processing). We evaluated the performance of Fast-Higashi and baselines under various evaluation metrics including: (1) the modularity score, (2) the adjusted rand index (ARI) and adjusted mutual information (AMI) scores, and (3) the Micro-F1 and Macro-F1 scores (see **Methods** for details). We made direct comparisons of Fast-Higashi against three scHi-C embedding methods, including two very recently developed scHi-C embedding methods, Higashi [20] and 3DVI [19] as well as scHiCluster [18] (which has been updated recently). It has been suggested that the updated scHiCluster can distinguish neuron subtypes better on the Lee et al. [15] dataset while the earlier version of scHiCluster cannot achieve [15, 20].

As shown in **Fig**. 2a-c, the UMAP visualizations of the Fast-Higashi embeddings on these three datasets show clear clustering patterns consistently corresponding to the annotated cell types. Importantly, we observed several major advantages of the Fast-Higashi embeddings compared to other methods. On the Lee et al. dataset of the human prefrontal cortex, based on the UMAP visualization of the embedding results, Fast-Higashi can resolve the differences among neuron subtypes clearly, separating all Pvalb, Sst, Vip, Ndnf, L2-3, L4, L5, and L6 neuron subtypes and showing more detailed structures within some cell types (**Fig**. 2b, marked in red box). To the best of our knowledge, this is the first time that excitatory neurons of different layers can be separated by using chromatin interaction information only. As a comparison (**Fig**. S2), the embeddings from Higashi and scHiCluster, while separating most of the neuron subtypes, have much weaker capability to distinguish excitatory neurons of different layers. The embeddings of 3DVI separate neurons into two categories, excitatory neurons and inhibitory neurons, lacking the ability for more refined cell type delineation. Moreover, the embeddings from both scHiCluster and 3DVI show obvious batch effects (**Fig**. S2).

**Figure 2:**
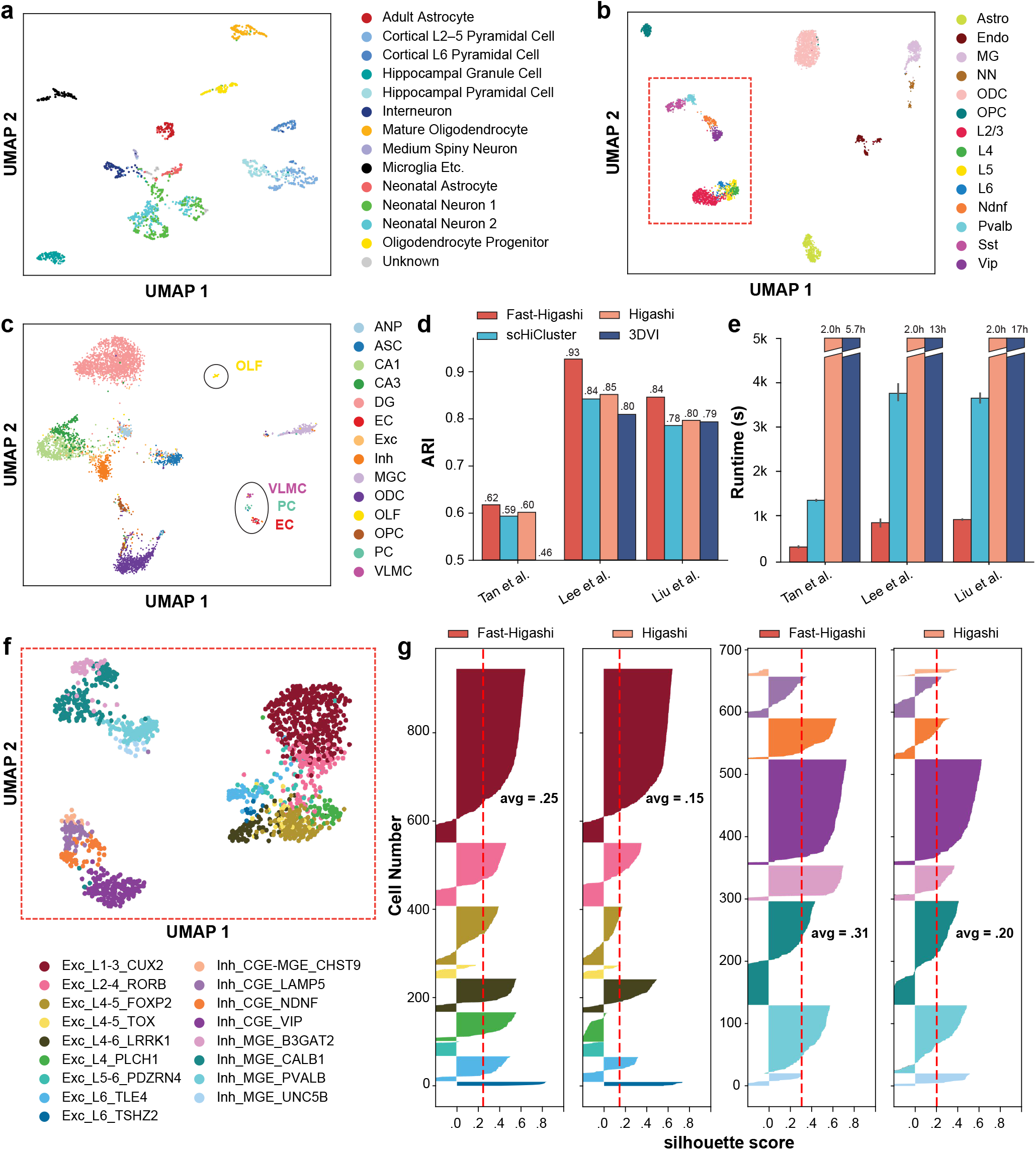
Evaluation of Fast-Higashi for generating embeddings for scHi-C data. **a**. UMAP visualization of the Fast-Higashi embeddings for the Tan et al. dataset [14]. See also **Fig**. S9. **b**. UMAP visualization of the Fast-Higashi embeddings for the Lee et al. dataset [15]. Cells in the red box are neuron cells. **c**. UMAP visualization of the Fast-Higashi embeddings for the Liu et al. dataset [16]. See also **Fig**. S3. **d**. Quantitative evaluation based on adjusted rand index (ARI) scores of the Louvain clustering results for each scHi-C embedding methods. See also **Fig**. S4. **e**. Runtime of different embedding methods across different datasets. **f**. UMAP visualization of the Fast-Higashi embeddings for the neuron cells in the Lee et al. dataset (cells in the red box in **(b)**). Cell type information is from Luo et al. [23]. See also **Fig**. S2. **g**. Quality of the embeddings for the neuron cells in the Lee et al. dataset measured as silhouette coefficients for neuron subtypes. See also **Fig**. S8. All cell type abbreviations are consistent with the data source.

On the Liu et al. dataset of the mouse hippocampus, we again observed Fast-Higashi’s clear advantage over other methods. Fast-Higashi is the only method that can separate CA3 cells from CA1 cells. More importantly, it can successfully identify small clusters of VLMC, PC, and EC cells, while all other methods except Higashi cannot (**Fig**. 2c and **Fig**. S3).

All these observations are supported by our quantitative results, where Fast-Higashi consistently achieves the highest or second best scores across all metrics of all three datasets (**Fig**. 2d and **Fig**. S4). We repeated the evaluation on two sci-Hi-C datasets with relatively lower coverage and/or a smaller number of cells and reached similar conclusions (see **Supplementary Results** B.2 for details).

In addition, we assessed the runtime of all scHi-C embedding methods. As shown in **Fig**. 2e, Fast-Higashi is significantly faster than all existing scHi-C embedding methods, especially the neural-network based methods (*>*40x faster than 3DVI and *>*9x faster than Higashi on the scHi-C datasets used for benchmarking). The runtime of Higashi mostly depends on its number of training epochs and is almost constant for datasets with more than 1000 cells.

Together, these results demonstrate that Fast-Higashi achieves the state-of-the-art performance for scHi-C embeddings with an ultrafast computational efficiency.

### Fast-Higashi enables the identification of rare cell types in complex tissues

In addition to the global evaluation on how well the Fast-Higashi embeddings correspond to the annotated cell types from the original datasets, we also sought to demonstrate that Fast-Higashi has unique capabilities to further improve the annotation of rare cell types in complex tissues.

We first visualized the Fast-Higashi embeddings of the neuron cells in the Lee et al. dataset using the UMAP projection (**Fig**. 2f). We obtained a new cell type annotation from [23], where the methylation profiles of the Lee et al. dataset were jointly embedded with single-cell methylation profiles from snmC-seq, snmCT-seq, and snmC2T-seq on human PFC to annotate cell types. This joint analysis allows the characterization of neuron subtypes in the Lee et al. dataset at a much more refined resolution. Based on the UMAP visualization, we observed that the smaller clusters within the same cell type (red box in **Fig**. 2b) can in fact be delineated into more detailed cell subtypes. For instance, the two finer clusters of Sst in **Fig**. 2b correspond to the CALB1 and B3GAT2 cell subtypes in **Fig**. 2f. By comparing with the UMAP visualization of other embedding methods (**Fig**. S2), we found that Fast-Higashi has the best ability to distinguish neuron subtypes, especially for the excitatory neurons. For inhibitory neurons, both Fast-Higashi and Higashi perform well and are the only two methods that can identify a smaller cluster of the UNC5B type (**Fig**. S2). To further support these observations, we also evaluated each method’s ability of separating neuron subtypes on the Lee et al. dataset through silhouette score analysis (**Fig**. 2g and **Fig**. S8). Consistent with our observations based on the UMAP visualization, Fast-Higashi achieves the highest average silhouette score on the neuron subtypes.

We next systematically evaluated the robustness of Fast-Higashi’s ability of identifying rare cell types, by simulating scHi-C dataset with different coverage. Specifically, we downsampled the contact pairs from the Lee et al. dataset to 10% to 50% of the original dataset and applied Fast-Higashi and Higashi, two models with strongest performance on these simulated datasets. As demonstrated by the UMAP visualizations in **Fig**. S5a, Fast-Higashi is more robust to the coverage of the dataset than Higashi, showing clearer clustering patterns that correspond to the cell types. The quantitative evaluation (**Fig**. S5b) further supports this observation.

Additionally, we applied Fast-Higashi to the Tan et al. dataset of the developing mouse brain. Fast-Higashi is able to separate most of the cell types marked from the data source (**Fig**. 2a). As compared to the existing scHi-C embedding methods, Fast-Higashi preserves the local trajectory of the time course better (**Fig**. S9). Surprisingly, we found two small clusters within the interneuron cell types and two separate clusters of neonatal neurons that do not correspond to the original Neonatal Neuron 1/2 labels from the dataset. These patterns are absent from the embeddings of other existing embedding methods except Higashi (**Fig**. S9). With the observation on the Lee et al. dataset that the small clusters within a cell type could reflect more refined subtypes, we believe that this could also be the case for the Tan et al. dataset. Detailed results will be discussed in a later section.

Taken together, these results confirm the unique capability of Fast-Higashi for identifying rare cell types or subtypes based on scHi-C data only.

### Fast-Higashi effectively captures cell-type specific 3D genome structures

We then sought to demonstrate that the meta-interactions captured by Fast-Higashi reflect the cell type-specific 3D genome features and can be used to interpret the generated embeddings. As a proof-of-principle, we first analyzed the meta-interactions of chromosome 1 for the Kim et al. dataset [12]. In this section, we mainly focused on the 4 cell types with enough cell numbers, including GM12878, H1ESC, HAP1, and HFFc6. We first visualized the single cell loadings of these meta-interactions (chromosome-specific embeddings). As shown in **Fig**. 3a, each cell type has its preferred set of meta-interactions. Note that due to the utilization of SVD (singular value decomposition) for solving the meta-interaction during the optimization process (see **Supplementary Methods** for details), the first meta-interaction (sorted by the singular value during the SVD process) would correspond to the general contact patterns of all cells within the scHi-C data. This is consistent with the observation that for all cells, their loadings of the first meta-interaction (marked as “1st MI” in **Fig**. 3a) are large and similar across all cell types. For all other meta-interactions, they represent how the cell type-specific 3D genome features deviate from the population interaction patterns. To validate this, we aggregated the cell type-specific meta-interactions weighted by the average single cell loadings and made comparisons to the differential contact patterns calculated from the bulk Hi-C. Specifically, we first calculated a “common bulk Hi-C” as the average of the bulk Hi-C of the same four cell types. Then for each cell type we calculated the differential contact patterns as the difference between the bulk Hi-C of that cell type and the “common bulk Hi-C”. As shown in **Fig**. 3b, the differential contact patterns calculated using the bulk Hi-C share similar patterns to the aggregated cell type-specific meta-interactions. This observation is consistent with the phenomenon that the Spearman correlations between cell type specific meta-interactions and the corresponding differential contact map is the highest (**Fig**. 3c).

**Figure 3:**
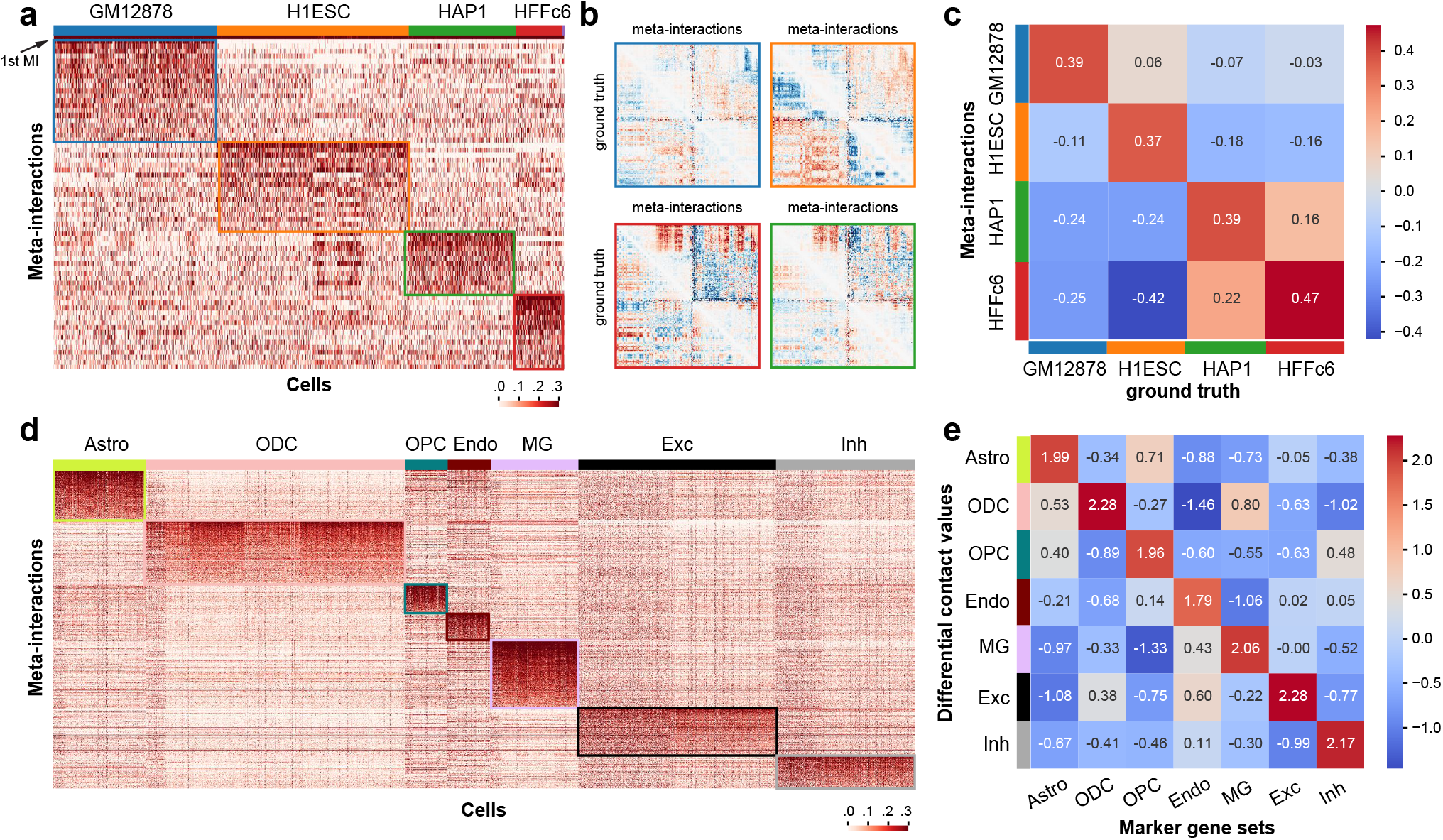
Analysis of the chromatin meta-interactions generated by Fast-Higashi. **a**. Heatmap of the single cell loadings for each meta-interaction of chromosome 1 for the Kim et al. dataset [12]. **b**. Visualization of the differential contact maps generated based on meta-interactions and those generated based on bulk Hi-C (marked with “ground truth”). Border color matches the cell type color in **(a). c**. Spearman correlation between differential contact maps generated based on meta-interactions and those generated based on bulk Hi-C (marked with “ground truth”). **d**. Heatmap of the single cell loadings for each meta-interaction of the whole genome for the Lee et al. dataset [15]. **e**. Mean differential contact values of the lists of cell type marker genes averaged for each cell type. The mean differential contact values are calculated using the corresponding meta-interactions as the summation of values for each bin in the meta-interaction contact map. For each cell type, the top 200 marker genes were identified using Seurat [24].

Next we analyzed the meta-interactions of a scHi-C dataset on complex tissues, i.e., the Lee et al. dataset [15]. **Fig**. 3d shows the single cell loadings of the whole genome meta-interactions for this dataset. We again can observe a clear preference of meta-interaction sets for different cell types. To confirm that these cell type-specific meta-interactions manifest cell type-specific 3D genome features that are functionally relevant, we calculated the differential contact values for each bin given a specific set of meta-interactions. We first aggregated the meta-interactions for a specific cell type by the average single cell loadings, leading to one meta-interaction map of size *N* × *N* for one cell type. We then calculated the differential contact values by summing over the column of this meta-interaction map, representing the overall deviation of a genomic bin from its population-level pattern. By comparing to the marker genes called from scRNA-seq [25, 26], we found that there is a strong positive correlation between the differential contact values and the expressions of the marker genes (**Fig**. 3e).

These results demonstrate that the meta-interactions from Fast-Higashi effectively capture the cell type-specific 3D genome features that are relevant to cell type-specific gene regulation. The meta-interactions from Fast-Higashi can be used to associate the embedding results to a specific region of the scHi-C contact map, pointing to further investigation of differential 3D chromatin contact patterns of various cell types in complex tissues.

### Fast-Higashi unveils single-cell 3D genome features in developing mouse brain

As discussed above, we applied Fast-Higashi to a scHi-C dataset of developing mouse brain, i.e., the Tan et al. dataset [14], and observed local clusters of cells within the two annotated cell types in the UMAP visualization (**Fig**. 2a). We postulated that these local clusters could potentially be subtypes not captured by other scHi-C embedding methods as well as the original data source. To demonstrate that Fast-Higashi can delineate finer scale cell types and uncover developmental trajectories, we first obtained Fast-Higashi embeddings for all cortex cells and annotated the observed small clusters as Interneuron (A), Interneuron (B), Neonatal Neuron (A), and Neonatal Neuron (B) (**Fig**. 4a).

**Figure 4:**
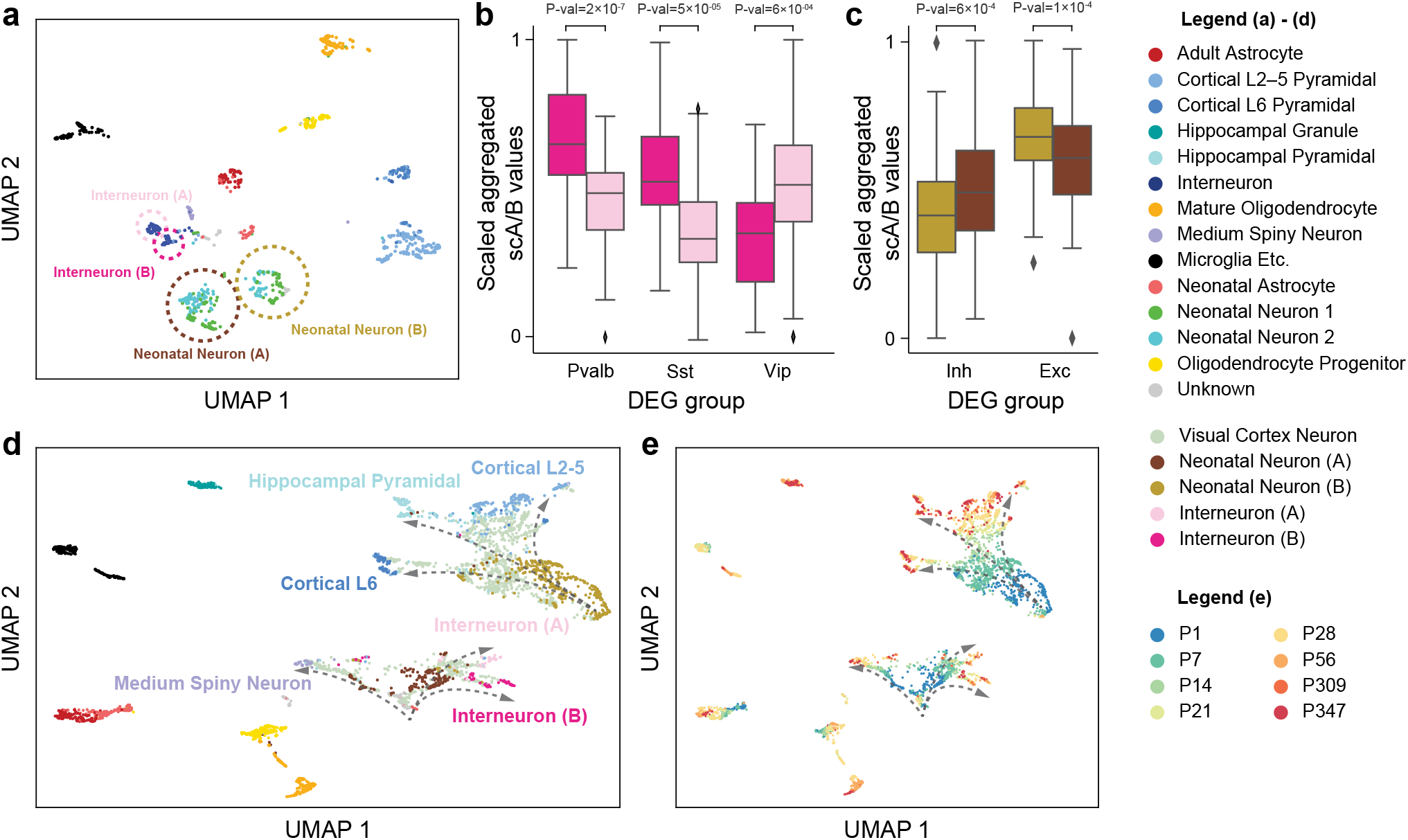
Application of Fast-Higashi to the scHi-C dataset from Tan et al. [14] of the mouse developing brain for more detailed identification of cell types and developmental trajectories. **a**. UMAP visualization of the Fast-Higashi embeddings for the cortex cells in the Tan et al. dataset. Cell subtypes identified by Fast-Higashi are highlighted with circles and texts. **b**. Distribution of the scaled aggregated single-cell A/B values for interneuron (A) and interneuron (B) subtypes identified by Fast-Higashi. For better visualization, the aggregated single-cell A/B values are linearly scaled to the range from 0 to 1 for each differentially expressed gene (DEG) group. **c**. Distribution of the scaled aggregated single-cell A/B values for Neonatal Neuron (A) and Neonatal Neuron (B) subtypes identified by Fast-Higashi. The scaling is the same as in panel **(b). d-e**. UMAP visualization of the joint Fast-Higashi embeddings of visual cortex, cortex, and hippocampus scHi-C datasets in Tan et al [14]. The potential developmental trajectories from neonatal neurons to fully mature neurons are marked by dashed arrows. Note that here **(d)** is colored with cell type labels and **(e)** is colored with ages of the mouse.

We sought to confirm our refined cell type labels of neonatal neurons and interneurons (highlighted with circles in **Fig**. 4a) by scA/B values in the gene bodies of marker genes. The marker genes were obtained from [14], which were calculated using Seurat [24] on the MALBAC-DT of the developing mouse brain. We quantified the A/B compartments of a set of differentially expressed genes (DEGs) by the aggregated scA/B value (see **Supplementary Methods** A.3 for details.) Previous studies reported the existence of global correlations between the scA/B values and the gene expression level within the same cell type [27] and across different cell types in complex tissues [14, 20]. In particular, genes with higher scA/B values are more likely to be highly expressed. We found that the aggregated scA/B value of the marker genes of Pvalb and Sst is significantly higher in Interneuron (B) as compared to Interneuron (A) and the marker genes of Vip show the opposite trend (**Fig**. 4b). These results suggest that the DEGs of Pvalb and Sst neurons are expressed at a higher level in Interneuron (B) than in Interneuron (A) and that the DEGs of Vip neurons exhibit the opposite behavior, indicating that Interneuron (A) and (B) are more likely to be Vip and Pvalb/Sst, respectively. Similarly, the aggregated scA/B value of the marker genes of neonatal inhibitory neurons is higher in Neonatal Neuron (A) and the aggregated scA/B value of the marker genes of neonatal excitatory neurons is higher in Neonatal Neuron (B) (**Fig**. 4c), indicating the Neonatal Neuron (A) and (B) are neonatal inhibitory neurons and neonatal excitatory neurons, respectively. Importantly, the aggregated scA/B values of neonatal excitatory neurons do not show different distributions (*P>*0.2) between the Neonatal Neuron 1 and 2 in the original annotations from [14], confirming that our annotations are indeed a refinement. Collectively, we again demonstrate the advantage of Fast-Higashi in identifying finer cell types.

To further validate our refined subtype annotation, we jointly embed the main cortex and hippocampus dataset of Tan et al. with another dataset of the visual cortex of developing mouse brain [14]. The main cortex and hippocampus dataset consists of cells from mice at 6 ages: P1, P7, P28, P56, P309, and P347, while the visual cortex dataset includes cells of mice at ages P7, P14, P21, and P28, which covers the critical development period from P7 to P28 that was missed in the original main datasets. When we applied Fast-Higashi to the union of these datasets, it recovered the complex developmental trajectories of inhibitory neurons and excitatory neurons. Specifically, in the UMAP visualization (**Fig**. 4d), a part of cells from the visual cortex dataset (light green) connect Neonatal Neuron (A) to the 3 mature inhibitory neuronal types: Interneuron (A), Interneuron (B), and Medium Spiny Neuron. Similarly, a different set of visual cortex cells connect Neonatal Neuron (B) to the 3 excitatory neuronal types: Cortical L2-5, Cortical L6, and Hippocampal Pyramidal. Since the Neonatal Neuron (A) and Neonatal Neuron (B) are primarily composed of P1/7 cells, and the 6 mature neuronal types consist of almost only P28 or older cells, placing the P14∼28 cells between P1/7 cells and P28+ cells is consistent with the developmental process. Moreover, along the inferred developmental branches (**Fig**. 4e (curved arrows)), cells are indeed ordered by the mouse ages, strongly supporting the ability of Fast-Higashi in recovering trajectory from scHi-C datasets. As a comparison, we included the results of scHiCluster (see **Fig**. S10). Although cells are ordered by mouse age in the embedding space of scHiCluster, we found that inhibitory neurons and excitatory neurons are not separated and scHiCluster cannot delineate two distinct developmental trajectories of inhibitory neurons and excitatory neurons.

In summary, using the Tan et al. dataset [14], we have demonstrated the clear advantages of Fast-Higashi in unveiling finer cell types over existing methods as well as the unique ability of Fast-Higashi to characterize the cell-to-cell variability of 3D genome features along complex biological processes.

## Discussion

In this work, we developed Fast-Higashi, an ultrafast and interpretable framework for scHi-C data analysis. The novel generalization from core-PARAFAC2 to Fast-Higashi not only inherits its strong scalability, but also enables joint and interpretable modeling of meta-interactions and cell embeddings. The development and incorporation of the Partial RWR algorithm further improve the performance of Fast-Higashi with negligible impact to the scalability. Evaluations of Fast-Higashi using a wide range of real scHi-C datasets have demonstrated its effectiveness and scalability for inferring informative cell embeddings, enabling the delineation of rare cell types and the reconstruction of developmental trajectories. Besides, as a proof-of-principle, we identified cell type-specific meta-interactions that are related to cell type-specific gene transcription. Together, we have demonstrated the effectiveness, scalability, and interpretability of our new method Fast-Higashi.

Despite having superior effectiveness and interpretability for scHi-C analysis, especially for generating informative cell embeddings that facilitate rare cell type identification, it is important to note that Fast-Higashi is not developed to replace its predecessor Higashi [20]. For instance, Fast-Higashi uses a random-walk-with-restart based method for imputing the sparse contact maps, which is efficient but also has limited imputation power. As demonstrated in [20], the accurate imputation empowered by hypergraph representation learning is key to unveiling important 3D genome features related to cell type-specific gene regulation. On the other hand, the underlying relationship between the tensor representation and the hypergraph representation of scHi-C data makes Fast-Higashi in some way a quasi-linear version of Higashi and can thus be used to initialize the Higashi model. As a proof-of-principle, we found that the Fast-Higashi initialized Higashi model can indeed achieve even better performance than any of these two methods (**Fig**. S11).

Fast-Higashi can be further enhanced by incorporating multimodal single-cell omics data, such as single-cell RNA-seq data and single-cell methylome data. Jointly modeling of co-assayed scHi-C data and other multimodal data has the potential to further improve cell embeddings and to establish connections between different modalities. Fast-Higashi may also be applied to study DNA-RNA interactions in single cells [28].

The further development of scHi-C related technologies is expected to expand rapidly in the coming years. Fast-Higashi has the potential to become an essential method in the toolbox of single-cell epigenomic analysis to greatly enhance the integrative investigation of 3D genome organization, genome functions, and cellular phenotypes at single-cell resolution for a wide range of biological applications.

## Methods

The design of Fast-Higashi is based on a tensor decomposition model, called core-PARAFAC2 [29], and is generalized to simultaneously model multiple 3-way tensors that share only a single dimension (single cells). The core-PARAFAC2 model is usually used to analyze multimodal data where observations may not be aligned along one of its modes. A concrete example in other applications is the electronic health records that contain multimodal phenotypes of multiple patients at various time points. Because a particular disease stage may begin at different time points and may have varying lengths across patients, a critical difficulty is that it is hard to align observations of different patients along the temporal dimension. Similarly, in scHi-C contact maps, TAD-like structures usually have varying sizes and boundaries in different genomic bins, obscuring the direct alignment of genomic bins. Therefore, we have developed Fast-Higashi based on core-PARAFAC2 to address this issue. In the following sections, we first introduce how Fast-Higashi performs tensor decomposition on the scHi-C datasets assuming that contact maps of only one chromosome are present. We discuss next how we generalize to multi-chromosome cases. We then derive the optimization procedure and introduce the novel partial random walk with restart (Partial RWR) module to address the sparseness of single-cell Hi-C dataset efficiently.

### Problem formulation of the Fast-Higashi model

For a scHi-C dataset, let ***𝒞*** denote the set of chromosomes. We formulate a collection of scHi-C contact maps of chromosome *c* ∈ ***𝒞*** as a 3-way tensor, denoted by 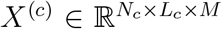, where *N*_*c*_ is the number of genomic loci (also denoted as genomic bins) in chromosome *c, L*_*c*_ is the number of features at each bin, and *M* is the number of cells in this dataset. In principle, *L*_*c*_ need not be equal to *N*_*c*_ because, for example, we may use different resolutions for genomic bins along the two dimensions and even include additional epigenomic features. However, for convenience, here we only consider contact maps and use the same resolution for both dimensions. We assume that *X*^(*c*)^ follows a 3-way core-PARAFAC2 model which includes: (1) a 3-way tensor 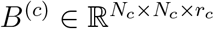 of *r*_*c*_ meta-interactions; (2) a matrix 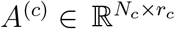 of bin weights indicating importance for each bin in every meta-interaction; (3) a chromosome-specific transformation matrix 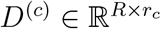; and (4) an orthogonal matrix *V* ∈ ℝ^*M ×R*^ that contains cell embeddings and is shared across all chromosomes, where *r*_*c*_ and *R* are hyperparameters.

We first introduce the cell-wise form of our model. As shown in **Fig**. 1a, the *𝓁*-th slice of *X*^(*c*)^ along the last dimension, denoted by 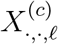, is the *𝓁*-th single-cell contact map, and we assume that it can be approximated by the weighted sum of meta-interactions:

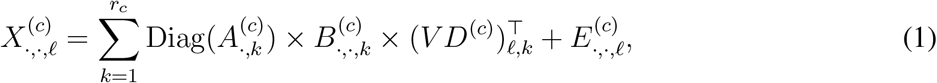

where 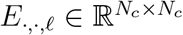 is a matrix of i.i.d. Gaussian noises with zero mean and arbitrary variance. Since *V* is the cell embedding matrix shared across all chromosomes, left multiplying *V* by the chromosome-specific transformation matrix *D*^(*c*)^ projects *V* to another space, which we term the chromosome-specific embedding space. The chromosome-specific embeddings *V D*^(*c*)^ directly quantify the contribution of each meta-interaction to single-cell contact maps, i.e., the overall weight of the *k*-th meta-interaction in cell *𝓁* is equal to (*V D*^(*c*)^)_*𝓁,k*_. Additionally, we also assume that bins in a meta-interaction may have different weights, i.e., the weight of bin *i* in the *k*-th meta-interaction is 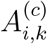. Together, the weight of the *k*-th meta-interaction at the *i*-th bin in cell *𝓁* is equal to the product of (1) the meta-interaction weight in the chromosome-specific embedding of cell *𝓁* and (2) the bin weight in the bin weight matrix, i.e. 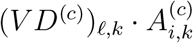.

To simplify the optimization problem, we introduce an alternative bin-wise form of this model [30]. Let 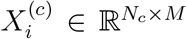 be the *i*-th slice along the first dimension of *X*^(*c*)^, i.e., the features of the *i*-th bin across all cells, and we use similar notations for other tensors. We assume that 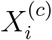 has the following decomposition:

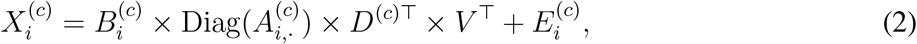

where 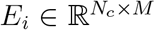 is a noise matrix. Since the noise is assumed to follow i.i.d. Gaussian distributions, the optimal set of parameters is the solution to the following optimization problem:

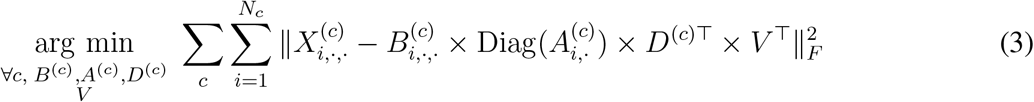

### Additional constraints for uniqueness

Now we introduce additional constraints to address the uniqueness issue of Eqn. 3 and to improve the ability of Fast-Higashi to capture critical topological patterns in single-cell Hi-C contact maps.

Without loss of generality, we show the uniqueness issue on a dataset with only one chromosome, denoted by *c*. Let (*B*(*c*), *A*(*c*), *D*(*c*), *V*) be one optimal solution to Eqn. 3. Then for any 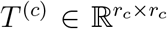 and 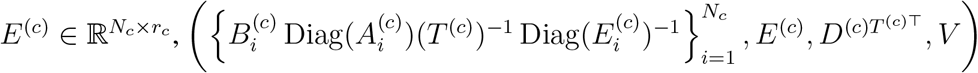 is also an optimal solution to Eqn. 3, implying the non-uniqueness. To address this, we impose constraints on the Gram matrices of 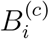 that,

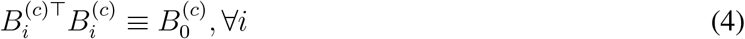

where 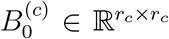 is constant over *i*. This is equivalent to requiring that 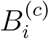 can be transformed to each other by left multiplying an orthogonal matrix, i.e., a rotation along the feature dimension.

These constraints enable Fast-Higashi to be less prune to noise and allow Fast-Higashi capture critical topological patterns from single-cell Hi-C contact maps. A concrete example is the TAD-like structure where the number of interactions within this region is expected to be higher but also non-uniform, in that, the near-diagonal elements usually include more interactions. The characteristics of a TAD-like structure cause the boundaries of this TAD-like structure to be the same for all bins in it but cause the location of the peak to vary across these bins. This indicates that it is impossible to directly find a pattern that fits more than one bin in this TAD-like structure. However, since we allow bin-specific rotations along the feature dimension, these rotations potentially can keep the boundary unchanged and redistribute the contacts among the features of each bin, allowing the shift of the peak. Matrices *A*^(*c*)^ are designed to capture the other bin-to-bin variability in scHi-C datasets. For example, bins usually have varying accessibility, which leads to different row sums in single-cell contact maps, and even in bins from one TAD-like structure. This variability is expected to be biologically meaningful and cannot be corrected by normalization. In Fast-Higashi, the bin weight matrix *A*^(*c*)^ will capture this variability. In addition, a single bin may also show cell type-specific accessibility, which is expected to be reflected as variation across meta-interactions in Fast-Higashi. In Fast-Higashi, these bin-specific and cell type-specific characteristics will be captured in the bin weight matrix *A*^(*c*)^ so that (1) any two bins may have different scaling factors in one meta-interaction and (2) one bin may have different scaling factors in any two meta-interactions. Therefore, these constraints retain the ability of capturing critical structures but also reduce the parameter spaces of Fast-Higashi, making it more robust to tolerate noise.

#### Algorithm 1

Optimization procedure for Fast-Higashi

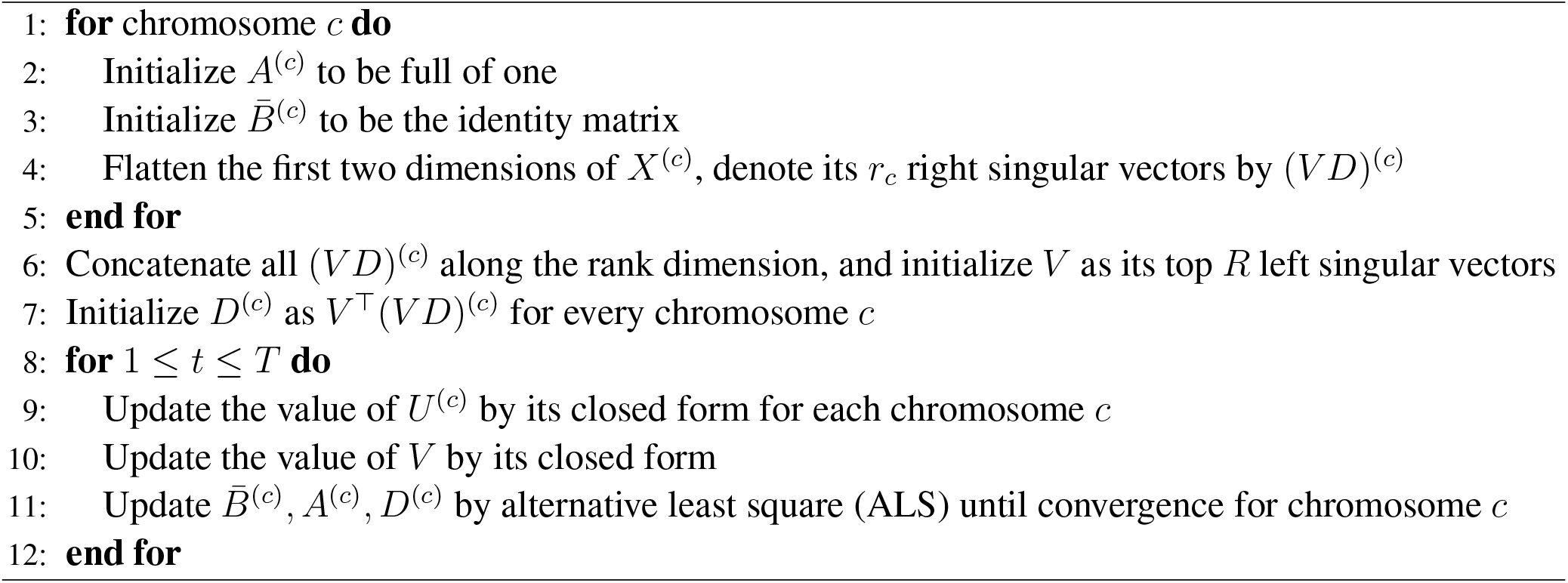

### Efficient parameter inference in Fast-Higashi

Here we show key steps in the derivation of a coordinate descent optimization procedure (summarized in Algorithm 1) for the optimization problem in Eqn. 3.

Since the derivations for different model parameters are similar, we only show the derivation of *V* as an example here. The complete derivation can be found in **Supplementary Methods**. For simplicity, let 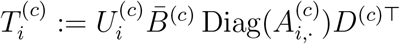, and then the optimization problem for *V* can be simplified as:

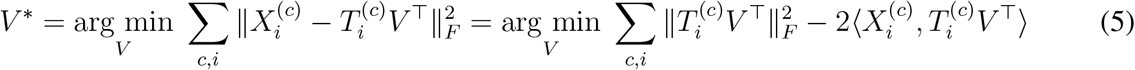

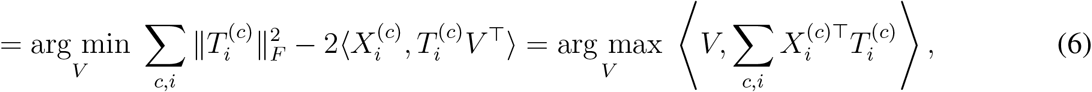

where the first equality in Eqn. 6 holds because ‖*TV* ^*T*^ ‖ _*F*_ = ‖*T* ‖_*F*_ for any orthogonal matrix *V*. Since *V* is orthogonal, the solution to this optimization has a closed form. Specifically, let the singular value decomposition (SVD) of 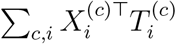 be 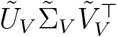 and the optimal solution of *V* is 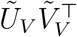.

### Initialization of the Fast-Higashi model

Here we provide efficient initialization of model parameters based on their interpretations (Algorithm 1 in **Supplementary Methods**). We initialize the matrix *A*^(*c*)^ to be full of one and the square matrix 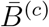 to be the identity matrix, for each chromosome. For chromosome *c*, we find the SVD of the single-cell contact maps of chromosome *c* and keep the top *r*_*c*_ right singular vectors which are the initial cell embeddings of chromosome *c*. To aggregate information from multiple chromosomes, we concatenate the initial cell embeddings from all chromosomes and find its SVD. We initialize the meta embedding *V* to be one of the orthogonal matrix and *D*^(*c*)^’s to contain the rest components in the SVD.

### Embedded partial random walk with restart (Partial RWR)

To mitigate the sparseness of the scHi-C contact maps, we sought to incorporate the random walk with restart (RWR) data imputation method [18] into the Fast-Higashi framework. However, direct utilization of RWR before the tensor decomposition process is not desirable. The RWR imputed contact maps are usually much denser than the original contact map, leading to much higher memory consumption for storing the results and lower computational efficiency for transforming data format between sparse matrices to dense tensors as well as data transferring between GPU and CPU. Our new solution is to integrate the RWR process during the optimization process of tensor decomposition and compute the RWR imputation batch by batch. The challenge for this design is that, as mentioned in the above section, the batch of the tensor decomposition optimization process is defined at the frontal slice of the tensor (Eqn. 3), i.e., the genomic bins, while the normal RWR requires the input of a complete graph adjacency matrix. To utilize RWR in our framework, here we propose the new partial random walk with restart (Partial RWR) algorithm. The procedures of this algorithm are shown in **Fig**. 1b, which consists of the following steps: For simplicity, in this section, we use *X* ∈ ℝ^*N ×N ×M*^ to represent the tensor representation of scHi-C contact maps of one chromosome. First, we fetch a small batch of the tensor *x*(*i*) := *X*_*i*:*i*+*bs*_ ∈ ℝ^*bs×N ×M*^ along the first dimension, where *bs* represents the batch size. Then, based on this small batch of tensor, we calculate the local affinity matrix *a*(*i, 𝓁*) ∈ ℝ^*bs×bs*^ of bins within this batch for each cell *𝓁*:

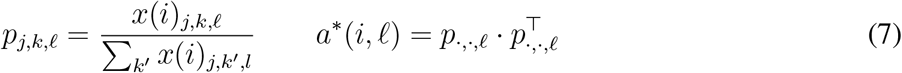

After that, the standard RWR algorithm is applied to these local affinity matrices:

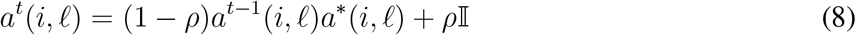

where *a*^0^(*i, 𝓁*) = 𝕀, and *ρ* is the restart probability in the RWR algorithm. We denote the converged results of the RWR algorithm as *a*^*∞*^(*i, 𝓁*) ∈ ℝ^*bs×bs*^ and use it as the weight to propagate the information from the original batch of the tensor *x*(*i, 𝓁*)

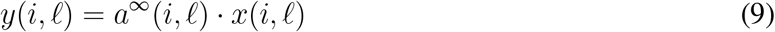

Finally, we use *y*(*i, 𝓁*) as the imputed results and pass it to the tensor decomposition optimization procedure. Our analysis showed that partial RWR can approximate the imputation of standard RWR well even with small batch sizes (see **Supplementary Results** B.1 for details). In this work, we use batch size 64 to keep the balance between accuracy and computational efficiency.

### Benchmarking scHi-C embedding methods

In this work, we mainly compared Fast-Higashi against three existing scHi-C embedding methods: Higashi [20], scHiCluster [18], and 3DVI [19] in terms of the quality of the generated embeddings and the runtime. We kept the embedding dimensions as the recommended ones for each method. For methods that allow selecting the maximum genomic distance to be considered, we set it to be 100Mb for all methods. We evaluated the embeddings generated by different methods under various evaluation metrics including: (1) Modularity score between the generated embeddings and the reference cell type label, (2) Adjusted rand index (ARI) and adjusted mutual information (AMI) score between the louvain clustering results and reference cell type label. Because the embeddings from different methods may reach the best clustering results at different combinations of parameters of Louvain clustering, we did a grid search for the number of neighbors and resolution parameters of the Louvain clustering for each method. The top 5 best clustering results for each method were kept and averaged as the final results. (3) We trained a logistic regression model using 10% of the cells and predicted the cell type for the rest 90% of cells. The Micro-F1 and Macro-F1 scores between the predicted cell type and reference ones were used to quantify the performance.

For the runtime analysis, all methods require different input formats and methods including 3DVI and Higashi can choose to only generate embeddings skipping the process of imputing sparse contact maps. To make a fair comparison, the runtime of all methods was calculated without the time of data processing, including transforming the scHi-C data into the format of a hypergraph in Higashi and reformatting the sparse contact maps into bands in 3DVI. For 3DVI and Higashi, we turned off the imputation function in the program and only used them to generate embeddings. For all methods, we added multiprocessing when possible even the multiprocessing was not originally implemented in some of the methods. Specifically, we parallelized the linear convolution and the random walk with restart algorithm of scHiCluster across all cells. We also parallelized 3DVI across different chromosomes, allowing the program to make full utilization of the GPUs. All methods were tested on a Linux machine with 1 NVIDIA RTX 2080 Ti GPU card, a 16-core Intel Xeon Silver 4110 CPU, and 252GB memory. All methods were set to use GPU when supported.

## Code Availability

Source code of Fast-Higashi can be accessed at: https://github.com/ma-compbio/Fast-Higashi.

## Acknowledgements

This work was supported by the National Institutes of Health Common Fund 4D Nucleome Program grant UM1HG011593 (J.M.) and the National Institutes of Health grants R01HG007352 (J.M.) and R01HG012303 (J.M.). J.M. is additionally supported by a Guggenheim Fellowship from the John Simon Guggenheim Memorial Foundation.

## Author Contributions

Conceptualization, R.Z., T.Z., and J.M.; Methodology, R.Z., T.Z., and J.M.; Software, R.Z., and T.Z.; Investigation, R.Z., T.Z., and J.M.; Writing – Original Draft, R.Z., T.Z., and J.M.; Writing – Review & Editing, R.Z., T.Z., and J.M.; Funding Acquisition, J.M.

