## Supplemental Information for "Ultrafast and interpretable single-cell 3D genome analysis with Fast-Higashi"

#### A Supplementary Methods

##### A.1 Data processing

In this work, we used several publicly available single-cell Hi-C datasets. We refer to them as Lee et al. [1] (GEO: GSE130711), Liu et al. [2] (GEO: GSE156683), Tan et al. [3] (GEO: GSE162511), Ramani et al. [4] (GEO: GSE84920), and Kim et al. [5] (4DN Data Portal: 4DNES4D5MWEZ, 4DNESUE2NSGS, 4DNESIKGI39T, 4DNES1BK1RMQ, and 4DNESTVIP977).

For all datasets except the Kim et al. dataset, we downloaded the contact pairs file from the corresponding GEO repository and transformed them into sparse contact maps at a given resolution (1Mb for the Ramani et al. dataset, 500Kb for the Lee et al., Liu et al., and Tan et al. datasets). For the Kim et al. dataset, we downloaded the FASTQ files from 4DN data portal and used the recommended processing pipeline (https://github.com/VRam142/combinatorialHiC).

##### A.2 Model inference in Fast-Higashi

Here, we derive a coordinate descent optimization procedure (summarized in Algorithm 1 in the main text) for the optimization problem in Eqn. 3. We also introduce necessary tricks for a GPU-compatible algorithm.

###### *A.2.1 Reformulation of the model*

To simplify the optimization problem, we introduce an alternative form of this model similar to [6]. Let $X^{(c)}_{i,\cdot,\cdot} \in \mathbb{R}^{N_c \times M}$ be the $i$-th slice along the first dimension of $X^{(c)}$ and we assume it has the following decomposition:

$$X^{(c)}_{i,\cdot,\cdot} = B^{(c)}_{i,\cdot,\cdot} \times \mathrm{Diag}(A^{(c)}_{i,\cdot}) \times D^{(c)\top} \times V^\top + E^{(c)}_{i,\cdot,\cdot}, \tag{10}$$

where $E_{i,\cdot,\cdot} \in \mathbb{R}^{N_c \times M}$ is a matrix of i.i.d. Gaussian noises with zero mean and arbitrary variance. To guarantee the uniqueness of the solution, two additional constraints are introduced:

$$B^{(c)\top}_i B^{(c)}_i \equiv C^{(c)} \quad \forall i \in [N_c], \quad \text{and} \quad \|A^{(c)}_{\cdot,\ell}\|_2 = \|D^{(c)}_{\cdot,\ell}\|_2 \quad \forall \ell \in [r_1], \tag{11}$$

where $C$ is an arbitrary matrix that is not dependent on $i$. With these constraints, the solution is unique up to a permutation along the dimension of length $r_c$ and a co-rotation of $V$ and $D^{(c)}$, which leads to a set of uniquely determined chromatin meta-interactions and chromosome-specific embeddings.

For optimization purpose, we first reformulate $B^{(c)}_i$ such that:

$$B^{(c)}_i = U^{(c)}_i \bar{B}^{(c)}, U^{(c)} \in \mathbb{R}^{N_c \times N_c \times r_c}, \bar{B}^{(c)} \in \mathbb{R}^{r_c \times r_c} \tag{12}$$

The uniqueness constraints in Eqn. 11 then become:

$$U_i^{(c)\top} U_i^{(c)} \equiv \mathbb{I} \quad \forall i \in [N_c], \quad \text{and} \quad \|\bar{B}_{\cdot,\ell}^{(c)}\|_2 = \|A_{\cdot,\ell}^{(c)}\|_2 = \|D_{\cdot,\ell}^{(c)}\|_2 \quad \forall \ell \in [r_c]. \tag{13}$$

The relation between $C^{(c)}$ in Eqn. 11 and $\bar{B}^{(c)}$ is $C^{(c)} = \bar{B}^{(c)\top} \bar{B}^{(c)}$.

##### A.2.2 Derivation for the optimal solution of $U_i^{(c)}$ and $V^*$

Now we derive the optimal value of $U_i^{(c)}$ given the rest parameters. For the sake of simplicity, let $T_U$ be $\bar{B}^{(c)} \operatorname{Diag}(A_{i,\cdot}^{(c)})(VD^{(c)})^\top$ and the optimization of $U_i^{(c)}$ can be simplified in as follows:

$$U_i^{(c)*} := \arg\min_{U_i^{(c)}} \|X_i^{(c)} - U_i^{(c)} T_U\|_F^2 = \arg\min_{U_i^{(c)}} \|X_i^{(c)}\|_F^2 + \|U_i^{(c)} T_U\|_F^2 - 2\langle X_i^{(c)}, U_i^{(c)} T_U\rangle \tag{14}$$

$$= \arg\min_{U^{(c)}} \|X_i^{(c)}\|_f^2 + \|T_U^\top\|_f^2 - 2\langle U_i^{(c)}, X_i^{(c)} T_U^\top\rangle = \arg\max_{U^{(c)}} \langle U_i^{(c)}, X_i^{(c)} T_U\rangle, \tag{15}$$

where the second to last equality is true because $\|UT\|_F = \|T\|_F$ holds for any orthogonal matrix $U$. Since $U^{(c)}$ is orthogonal, the solution to this optimization has a closed form. Specifically, let the SVD of $X_i^{(c)} T_U^\top$ be $\tilde{U}_U \tilde{\Sigma}_U \tilde{V}_U^\top$, and the optimal solution of $U_i^{(c)*}$ is $\tilde{U}_U \tilde{V}_U^\top$.

The closed form of $V^*$ is derived in a similar way. Let $T_i^{(c)} := U_i^{(c)} \bar{B}^{(c)} \operatorname{Diag}(A_{i,\cdot}^{(c)}) D^{(c)\top}$ and then

$$V^* := \arg\min_V \sum_{c,i} \|X_i^{(c)} - T_i^{(c)} V^\top\|_F^2 = \arg\max_V \left\langle V, \sum_{c,i} X_i^{(c)\top} T_i^{(c)} \right\rangle, \tag{16}$$

which implies $V^* = \tilde{U}_V \tilde{V}_V^\top$ where $\tilde{U}_V \tilde{\Sigma}_V \tilde{V}_V^\top$ is the SVD of $\sum_{c,i} X_i^{(c)\top} T_i^{(c)}$.

##### A.2.3 Derivation for the optimal solution of $\bar{B}^{(c)}$, $A^{(c)}$, and $D^{(c)}$

Next, we derive the optimization of $\bar{B}^{(c)}$, $A^{(c)}$, and $D^{(c)}$. For simplicity, we denote $\bar{B}^{(c)} \operatorname{Diag}(A_{i,\cdot}^{(c)}) D^{(c)\top}$ by $T_i$. Using the same trick, we can simplify the optimization in the following way:

$$\arg\min_{\bar{B}^{(c)}, A^{(c)}, D^{(c)}} \sum_i \|X_i^{(c)} - U_i^{(c)} T_i V^\top\|_F^2 = \arg\min_{\bar{B}^{(c)}, A^{(c)}, D^{(c)}} \sum_i \|T_i - U_i^{(c)\top} X_i^{(c)} V\|_F^2 \tag{17}$$

If we stack the $r_c$-by-$R$ matrices $U_i^{(c)\top} X_i^{(c)} V$ to create a 3-way tensor $Y^{(c)} \in \mathbb{R}^{N_c \times r_c \times R}$, the optimization becomes:

$$\arg\min_{\bar{B}^{(c)}, A^{(c)}, D^{(c)}} \left\| \sum_{k=1}^{r_c} A_{\cdot,k}^{(c)} \otimes \bar{B}_{\cdot,k}^{(c)} \otimes D_{\cdot,k}^{(c)} - Y^{(c)} \right\|_F^2, \tag{18}$$

which is exactly the PARAFAC model and $\bar{B}^{(c)}$, $A^{(c)}$, and $D^{(c)}$ can be solved by alternative least square (ALS) [7].

##### A.2.4 Mini-batch optimization

To improve the scalability of the method, we implemented the optimization of $U^{(c)}$ in a batch-wise manner. For a typical scHi-C dataset of 10,000 cells, if we set the resolution to 500Kb, the 3-way dense tensor of chromosome 1 that is ready for tensor operations takes up to 6GB GPU RAM, which leaves inadequate RAM for subsequent computations on GPU. To utilize the computation power of GPU, we divide $X^{(c)}$ and $U^{(c)}$ into batches along the first dimension, and update all the slices of $U^{(c)}$ from this batch in parallel using GPU. To minimize the data transfer amount between CPU and GPU, we compute the $T_i^{(c)}$ for the optimization of $V$ in Eqn. 16 before we remove the copy of this batch from GPU. Since $r_c$ is much smaller than $N_c$, the tensor $T^{(c)}$ fits in the GPU. Besides, we store these 3-way tensors $X^{(c)}$ in the COO format and transfer each batch of slices into GPU in the form of sparse COO tensors, which minimizes the data transfer as well as CPU memory usage. Hence, our method is optimized for GPU with limited RAM and data transfer rate to utilize its computation power and accelerate the overall running time.

##### A.3 Aggregated single-cell A/B value

We developed the aggregated single-cell A/B value to collectively quantify the chromatin conformation at multiple loci in one cell. We calculated the scA/B value of every 500Kb genomic locus in single cells following the method proposed in [3]. We defined the scA/B value of one gene as the average of the scA/B values of the genomic loci spanned by that gene. Although Higashi software includes an algorithm to to calculate scA/B values based on its embeddings and imputations, to avoid potential analysis bias, we instead used the more orthogonal method from [3]. It is worth noting that in [3], by using the calculated scA/B values as embeddings, the observed refined clustering results in Fast-Higashi embeddings do not exist. To summarize the behavior of a group of genes, we defined the aggregated scA/B value of these genes in one cell as the average scA/B value across all genes in that cell. To systematically assess the differential expression of a group of genes between two sets of cells, we examined the difference in the distribution of aggregated scA/B value of these genes between the two sets of cells by t-test.

### B Supplementary Results

#### B.1 Evaluation for Partial RWR

To evaluate how the Partial RWR algorithm would perform for sparse data imputation compared to the original RWR algorithm, we tested the impact of batch size on chromosome 1 of the Lee et al. dataset [1] at 500Kb resolution. Specifically, we applied Partial RWR of different batch sizes and RWR on the sparse contact maps and then calculated Pearson and Spearman correlation scores between the imputed results from Partial RWR and standard RWR. As expected, both similarity measurements increase with the batch size (**Fig.** S1). We also observed that even for a relatively small batch size such as 32 or 64, the correlation score would already surpass 0.9.

#### B.2 Evaluation on sci-Hi-C datasets

To further demonstrate that Fast-Higashi can work with scHi-C datasets with relatively lower coverage with a simpler cell type composition, we applied it to two sci-Hi-C datasets at 1Mb resolution: the Ramani et al. dataset [4] and the Kim et al. dataset [5] (**Fig.** S6). On these two relatively simpler datasets, all four embedding methods performed well based on the quantitative evaluation (**Fig.** S4). In terms of identifying cell types with a relatively small number of cells, Fast-Higashi and Higashi perform the best for the Ramani et al. dataset, where the GM12878 cells form into a small cluster. 3DVI performs the best on the Kim et al. dataset, where the IMR90 cells are clustered into a corner of the HFFc6 cell cluster.

In addition, we assessed the runtime of all scHi-C embedding methods on these two datasets. As shown in **Fig.** S7, Fast-Higashi is still much faster than all existing scHi-C embedding methods.

### C Supplementary Figures

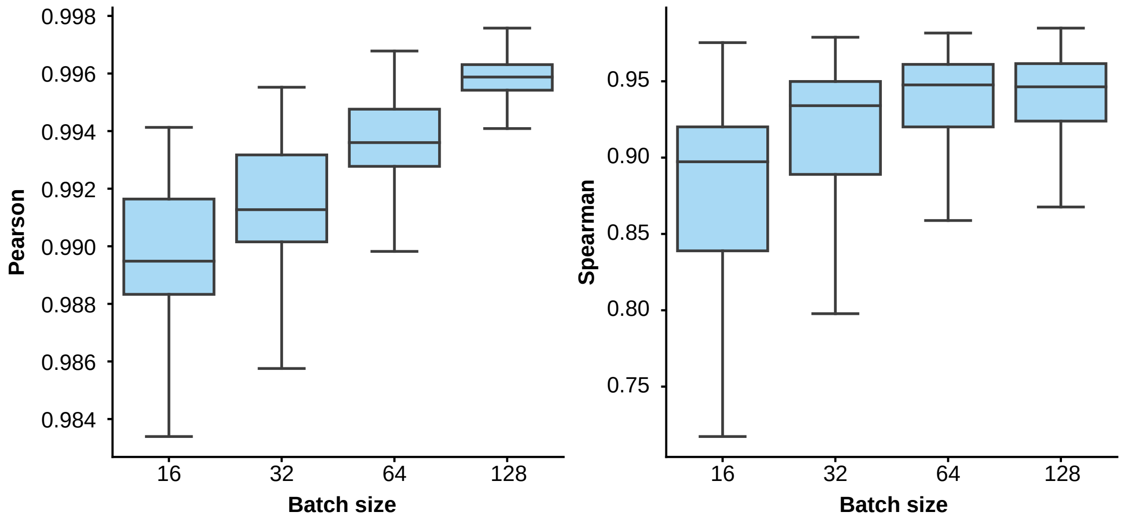


**Figure S1:** Similarity between the imputed results from Partial RWR and standard RWR with varying batch sizes.

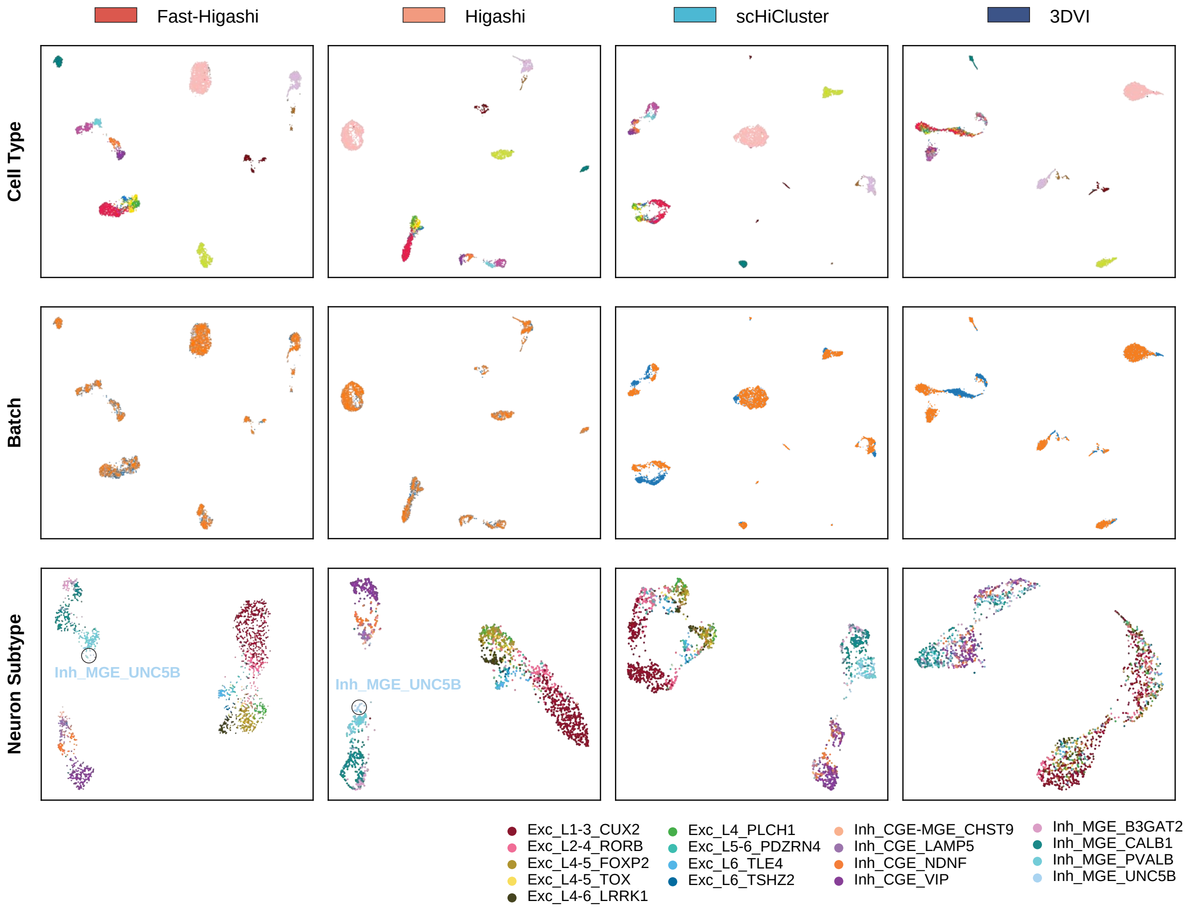


**Figure S2:** UMAP visualization of the embeddings from different scHi-C embedding methods for the Lee et al. dataset [1]. Scatter plots are colored with the cell type information from the original data source, the more refined cell type information from Luo et al. [8], and the batch id.

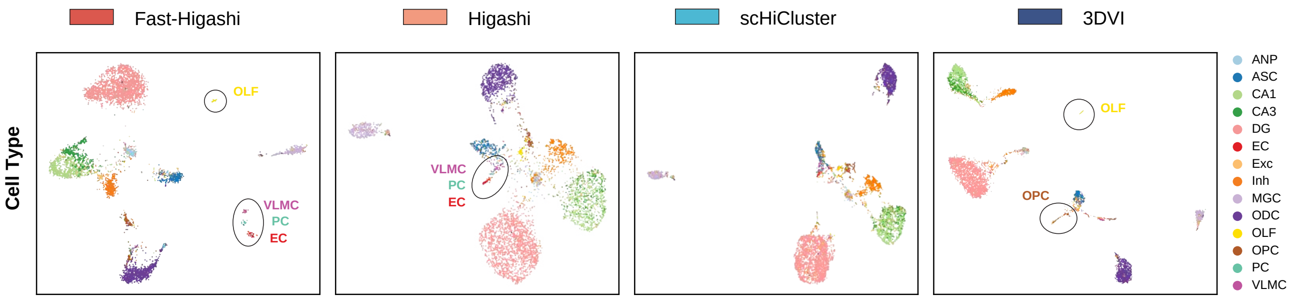


**Figure S3:** UMAP visualization of the embeddings from different scHi-C embedding methods for the Liu et al. dataset [2].

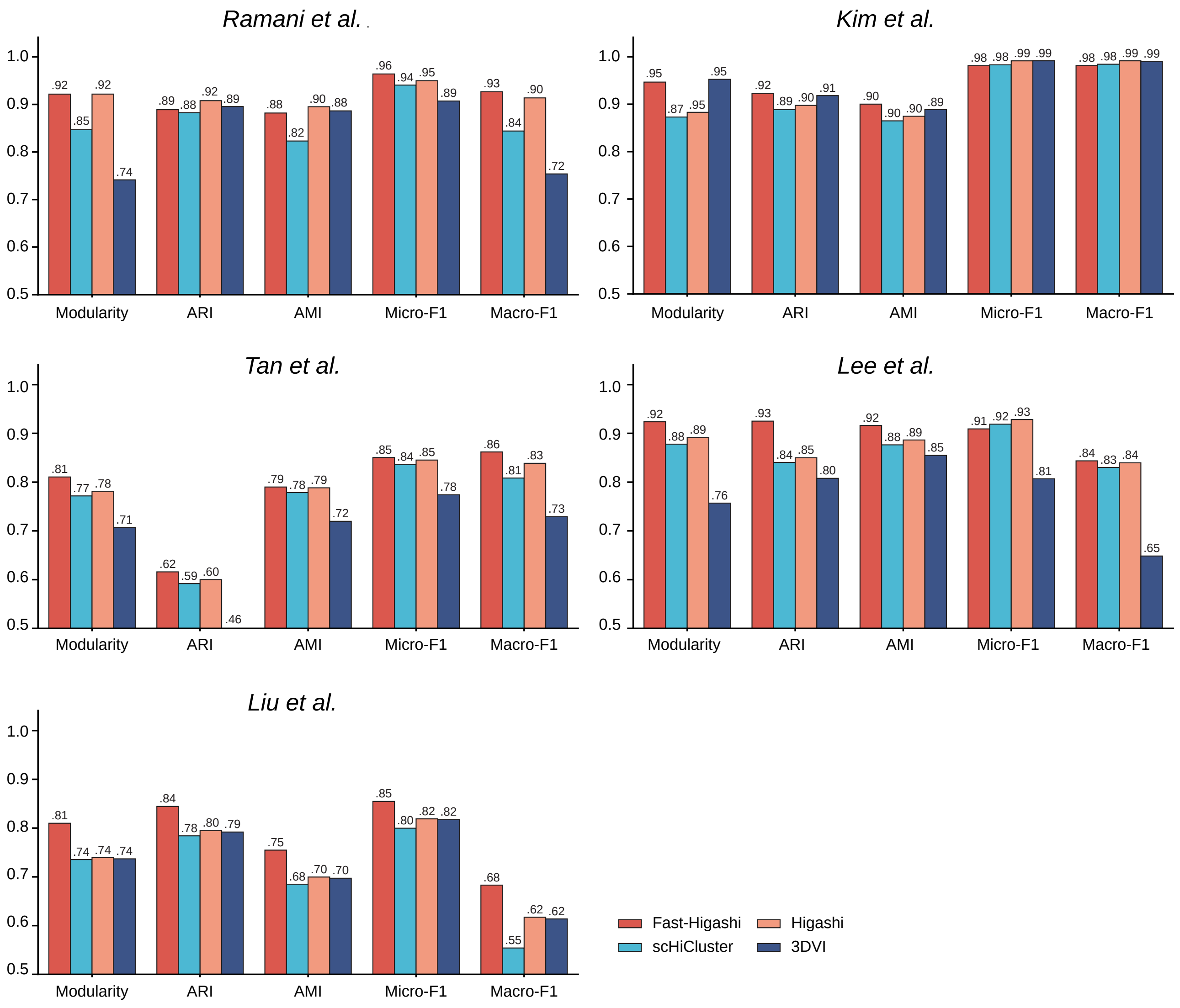


**Figure S4:** Quantitative evaluation for different scHi-C embeddings methods across different scHi-C datasets. The evaluation metrics include the modularity scores, similarity scores (adjusted rand index and adjusted mutual information) between the Louvain clustering results and the reference cell type label, and measurement of prediction accuracy (Micro-F1 and Macro-F1) between the predicted cell type and the reference cell type.

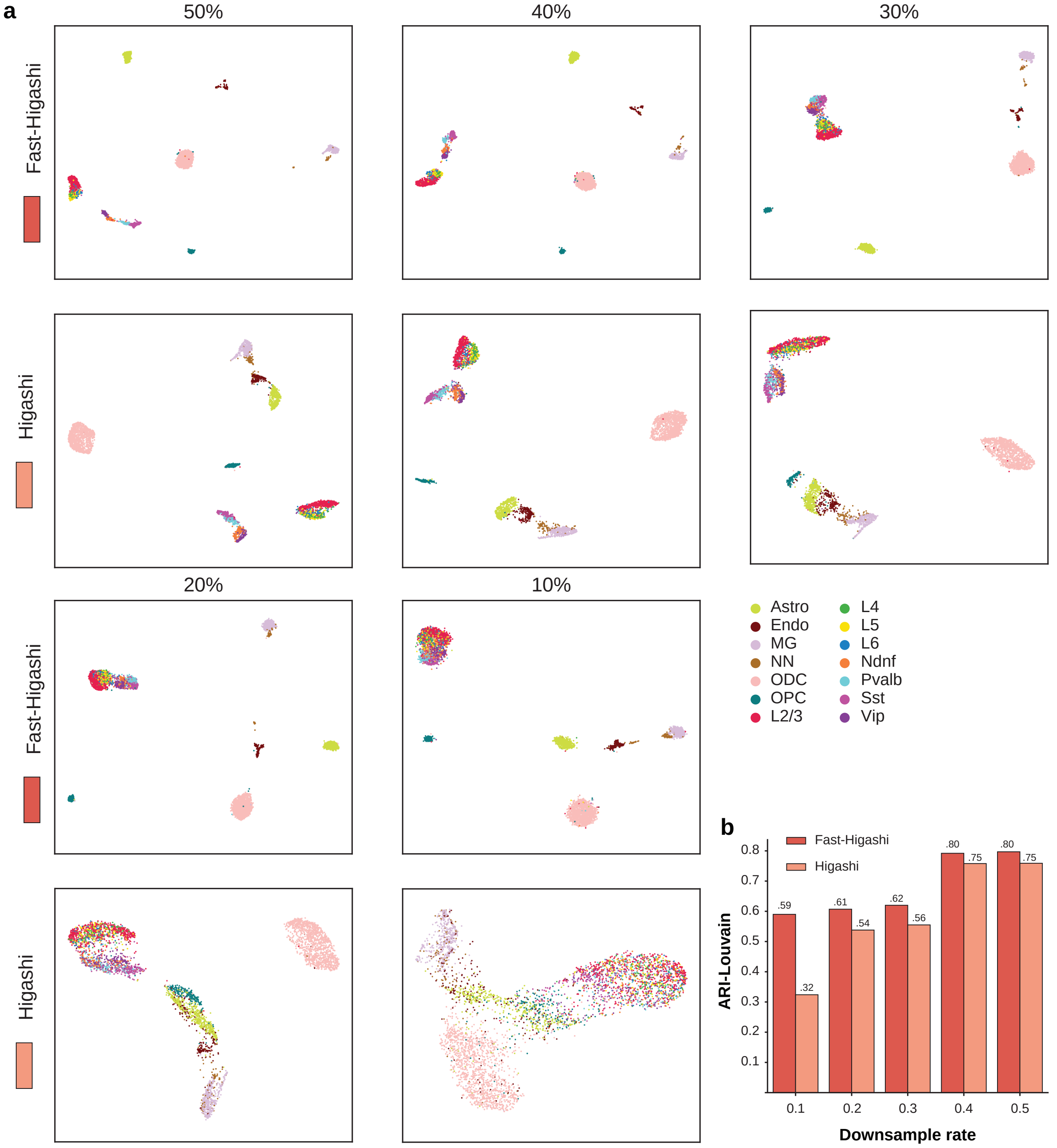


**Figure S5:** Comparisons of Higashi and Fast-Higashi embeddings for the Lee et al. dataset under different downsample ratio. **a.** UMAP visualization of embeddings from Higashi and Fast-Higashi for the Lee et al. dataset under different downsample ratio. b. Quantitative evaluation based on adjusted rand index (ARI) scores of the Louvain clustering results for Higashi and Fast-Higashi embeddings. The evaluation was carried out on the Lee et al. dataset with different downsample ratio

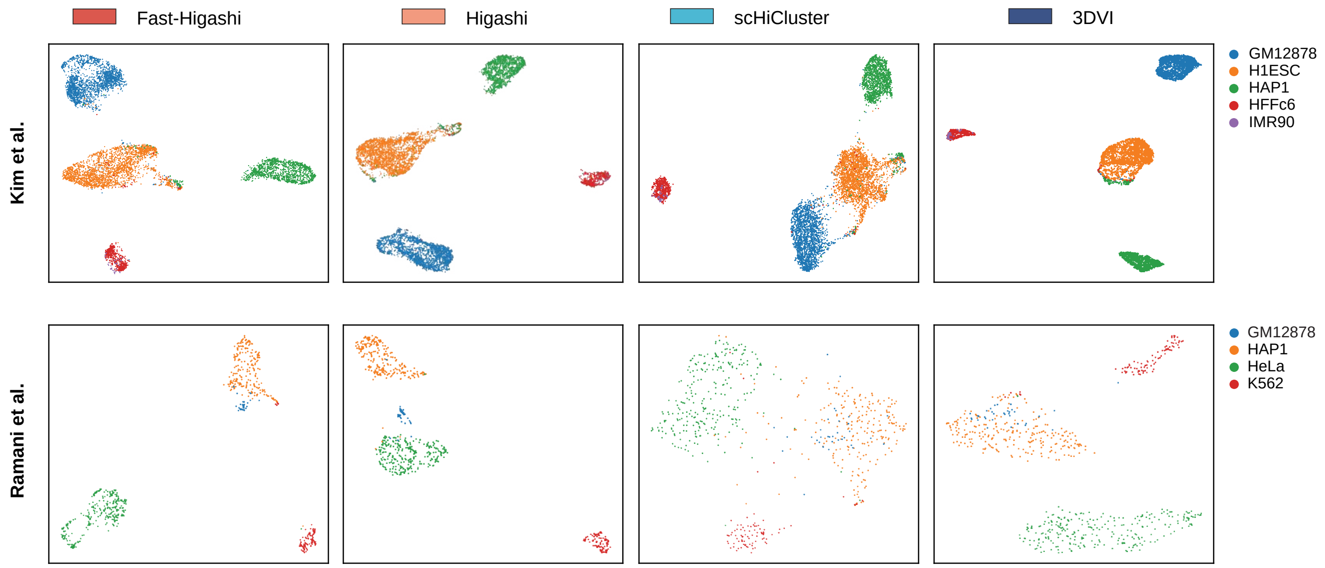


**Figure S6:** UMAP visualization of the embeddings from different scHi-C embedding methods for two sci-Hi-C datasets: the Ramani et al. dataset [4] and the Kim et al. dataset [5].

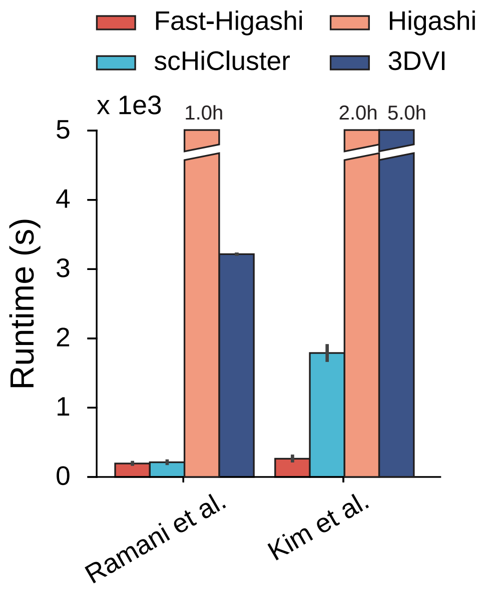


**Figure S7:** Runtime of different scHi-C embedding methods on two sci-Hi-C datasets.

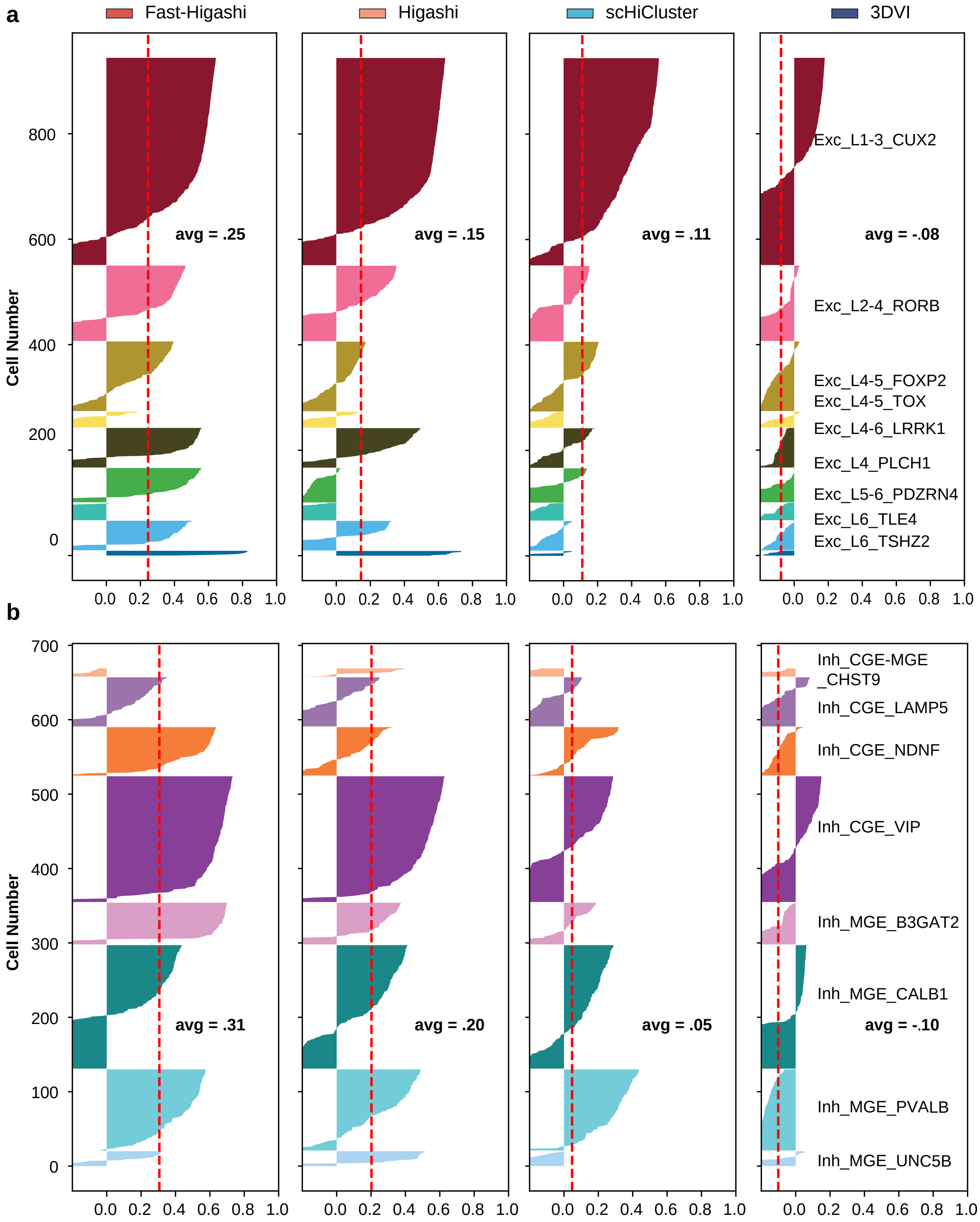


**Figure S8:** Quality of the embeddings for the neuron cells in the Lee et al. dataset measured as silhouette coefficients for neuron subtypes.

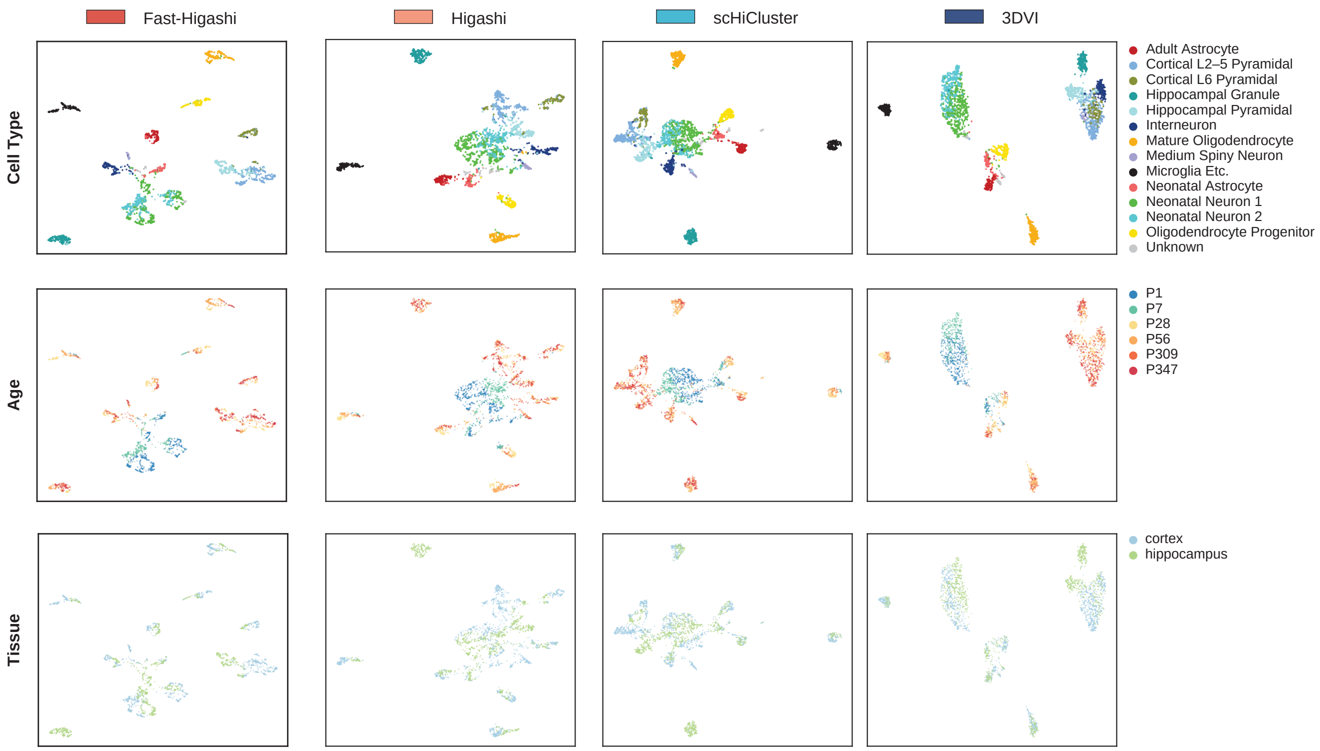


**Figure S9:** UMAP visualization of the embeddings from different scHi-C embedding methods for the Tan et al. dataset [3]. Scatter plots are colored with the cell type information, the ages of the mouse, and the tissue origin of the cells.

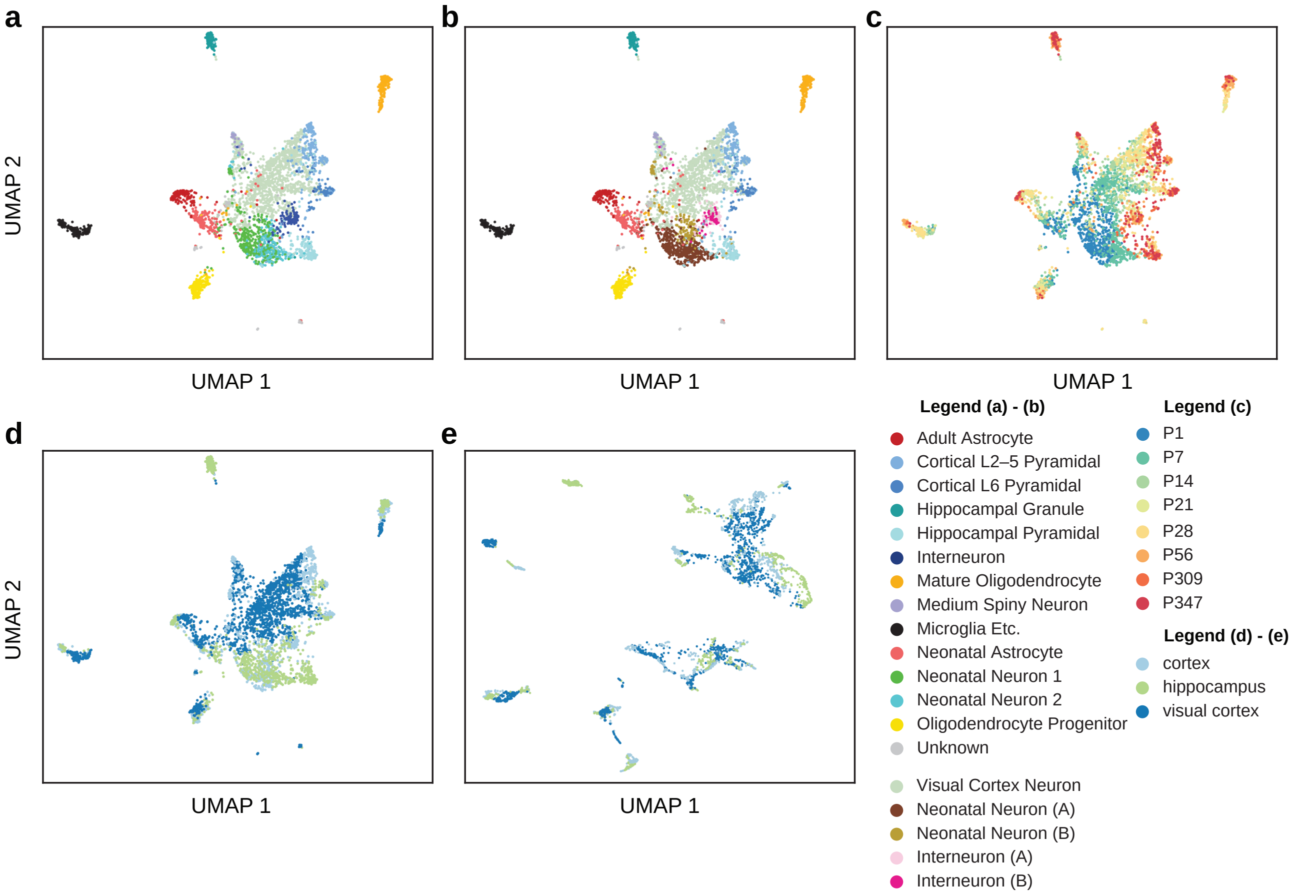


**Figure S10: a. b. c. d.** UMAP visualization of the joint embeddings from scHiCluster for visual cortex, hippocampus, and cortex tissues from the Tan et al. dataset. Colors reflect the original cell type information from Tan et al. [3], the refined cell type from this work, the ages of the mouse, and the tissue of origin of the cells, respectively. **e.** UMAP visualization of Fast-Higashi joint embeddings for visual cortex, hippocampus, and cortex tissues from the Tan et al. dataset (colored by the tissue of the cells).

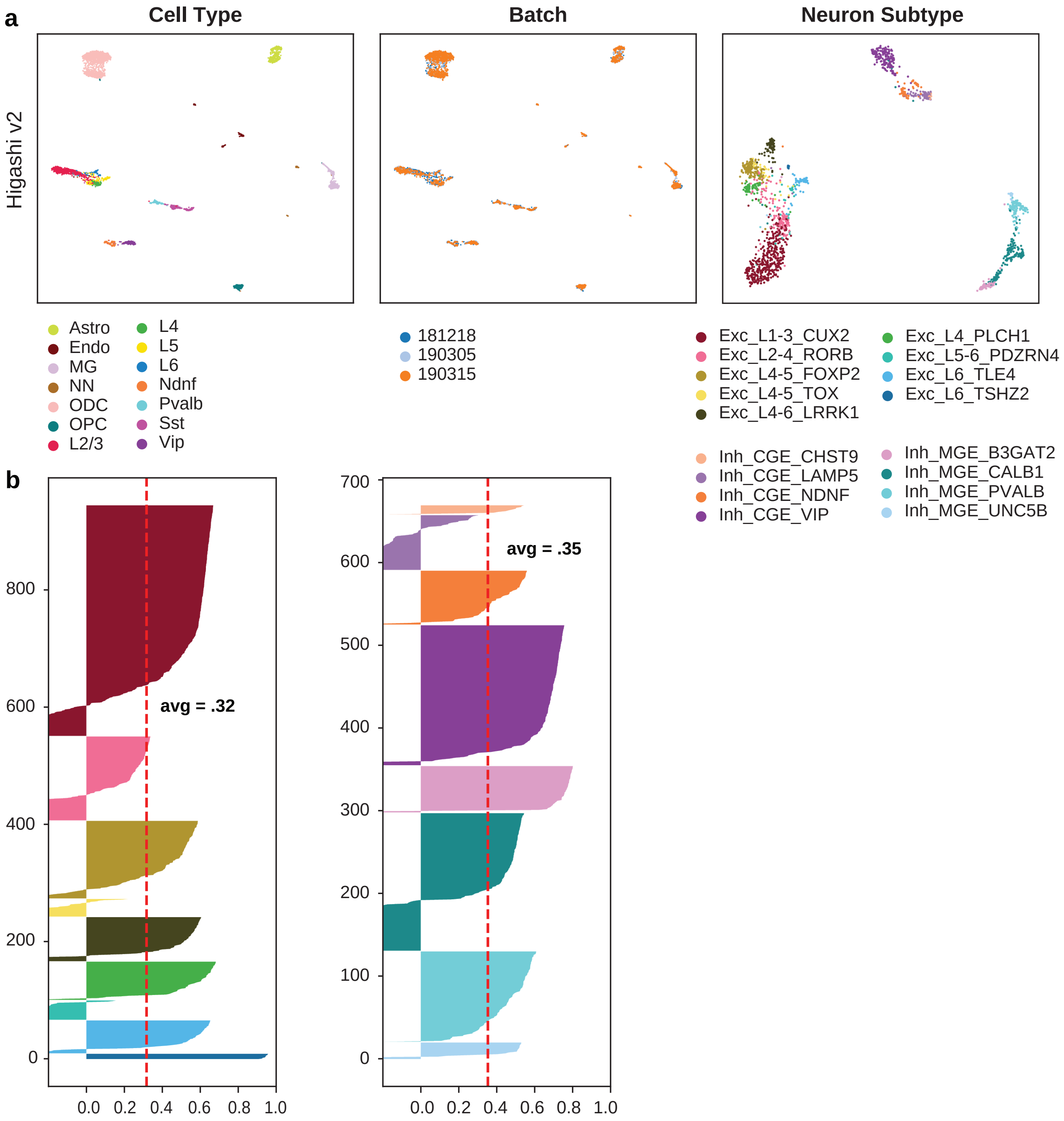


**Figure S11:** Embeddings from the Fast-Higashi initialized Higashi model on the Lee et al. dataset. **a.** UMAP visualization of embeddings. Scatter plots are colored with cell type information from the original data source, the more refined cell type information from Luo et al. [8], and the batch id. **b.** Quality of the embeddings for the neuron cells in the Lee et al. dataset measured as silhouette coefficients for neuron subtypes.
